# Temperature-dependent Spike-ACE2 interaction of Omicron subvariants is associated with viral transmission

**DOI:** 10.1101/2024.01.20.576353

**Authors:** Mehdi Benlarbi, Shilei Ding, Étienne Bélanger, Alexandra Tauzin, Raphaël Poujol, Halima Medjahed, Omar El Ferri, Yuxia Bo, Catherine Bourassa, Julie Hussin, Judith Fafard, Marzena Pazgier, Inès Levade, Cameron Abrams, Marceline Côté, Andrés Finzi

## Abstract

The continued evolution of SARS-CoV-2 requires persistent monitoring of its subvariants. Omicron subvariants are responsible for the vast majority of SARS-CoV-2 infections worldwide, with XBB and BA.2.86 sublineages representing more than 90% of circulating strains as of January 2024. In this study, we characterized the functional properties of Spike glycoproteins from BA.2.75, CH.1.1, DV.7.1, BA.4/5, BQ.1.1, XBB, XBB.1, XBB.1.16, XBB.1.5, FD.1.1, EG.5.1, HK.3 BA.2.86 and JN.1. We tested their capacity to evade plasma-mediated recognition and neutralization, ACE2 binding, their susceptibility to cold inactivation, Spike processing, as well as the impact of temperature on Spike-ACE2 interaction. We found that compared to the early wild-type (D614G) strain, most Omicron subvariants Spike glycoproteins evolved to escape recognition and neutralization by plasma from individuals who received a fifth dose of bivalent (BA.1 or BA.4/5) mRNA vaccine and improve ACE2 binding, particularly at low temperatures. Moreover, BA.2.86 had the best affinity for ACE2 at all temperatures tested. We found that Omicron subvariants Spike processing is associated with their susceptibility to cold inactivation. Intriguingly, we found that Spike-ACE2 binding at low temperature was significantly associated with growth rates of Omicron subvariants in humans. Overall, we report that Spikes from newly emerged Omicron subvariants are relatively more stable and resistant to plasma-mediated neutralization, present improved affinity for ACE2 which is associated, particularly at low temperatures, with their growth rates.

## INTRODUCTION

Since the beginning of the COVID-19 pandemic, multiple SARS-CoV-2 variants emerged, raising concerns about the efficacy of infection and/or vaccine-elicited immunity (1-5). A SARS-CoV-2 variant harboring 33 mutations in its Spike glycoprotein (S), Omicron (BA.1) reduced vaccine efficacy against infection due to its improved antibody escape (6-12). While modified versions of mRNA vaccines were produced to induce an immune response against the Omicron Spike (BA.1) (13, 14) and its subvariants (15-18), the persistent evolution of SARS-CoV-2 gave rise to various subvariants across the world (Figure 1A, 2A) (19). BA.1 was rapidly surpassed by BA.2 (20), and since then, several of its progenies have emerged and demonstrated improved transmission (21). Notably, BA.2.75, which surfaced in May 2022 further mutated into CH.1.1 (22, 23). On the other hand, BA.4 and BA.5 which harbor the same Spike further evolved to BQ.1.1 in late 2022, showing improved immune escape ability (24). In that same period, a recombinant sublineage, XBB, emerged and showed enhanced immune escape (25, 26). Since then, most newly occurring Omicron subvariants are derived from XBB and carry the Ser486Pro mutation known to enhance the affinity for the human receptor angiotensin converting enzyme 2 (ACE2) (27), such as XBB.1.5 which quickly dominated over XBB in January 2023 (28, 29). Recently, the main XBB subvariants were XBB.1.5, XBB.1.9.1, XBB.1.16, XBB.2.3 and EG.5.1, representing around 80% of reported viral sequences (30), in addition to EG.5.1 sublineages HK.3 and HV.1 which are growing rapidly around the globe (31, 32). In August 2023, the emergence of BA.2.86, a highly divergent BA.2 subvariant, caused great concerns regarding infection- and vaccine-elicited immune responses (30, 33, 34). Although the global number of infections related to BA.2.86 were relatively limited, its fast expansion and diversification in various countries has been noted, with the emergence of BA.2.86 sublineages (*i.e.,* JN.1, JN.2 and JN.3) showing enhanced transmission globally (32, 35). As of January 2024, JN.1 and its derivatives represent around 70% of reported viral sequences according to Nextstrain (https://nextstrain.org/ncov/gisaid/global/6m).

**Figure 1.**
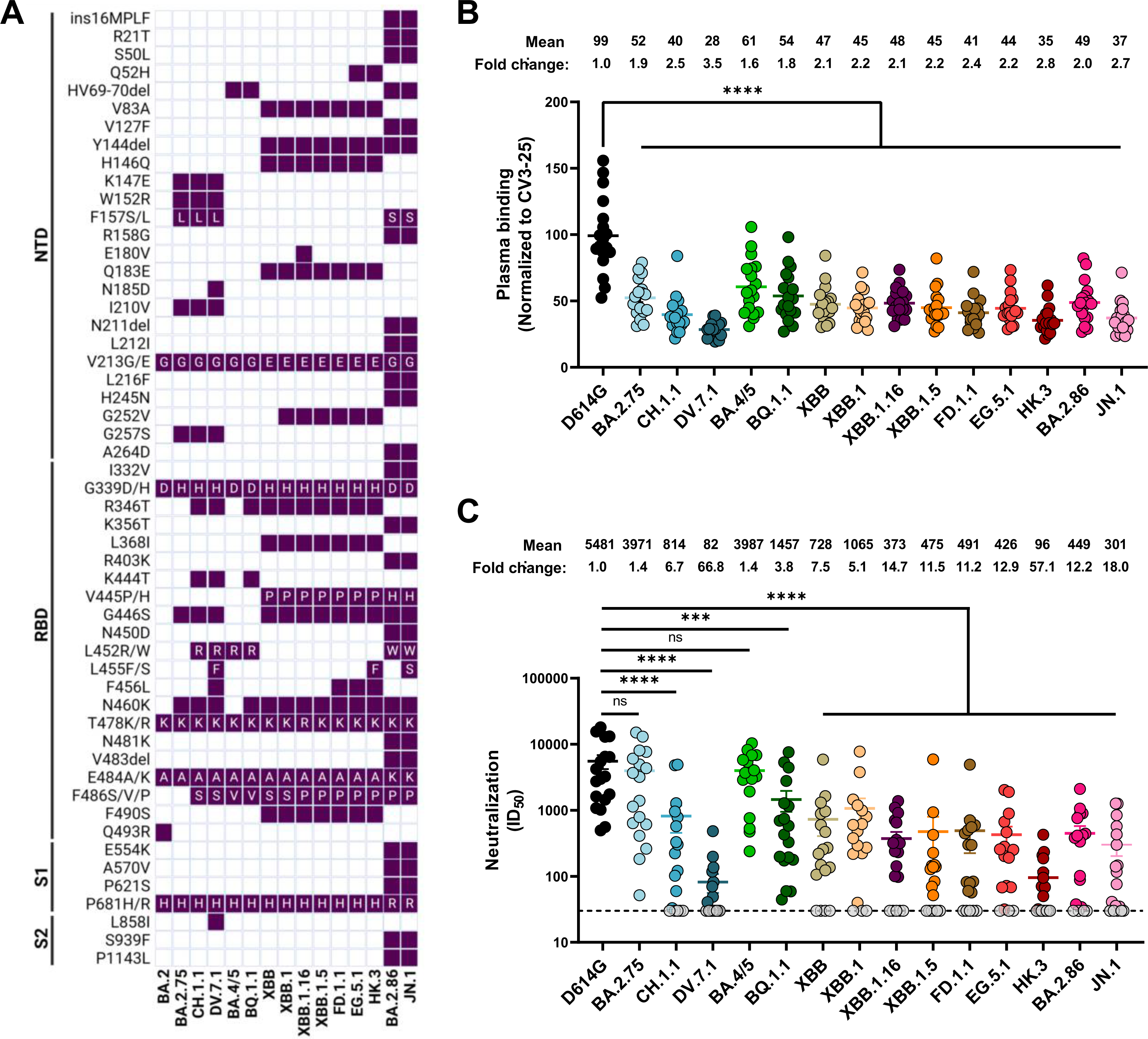
Plasma Recognition and Neutralization of emerging Omicron subvariants. (**A**) Defining Spike glycoprotein (S) mutations of recent Omicron subvariants in the N-terminal domain (NTD), receptor binding domain (RBD) and the rest of subunit 1 (S1) and subunit 2 (S2). (**B**) HEK293T cells were transfected with the indicated full-length S, stained with the CV3-25 mAb or with plasma from vaccinated individuals collected three weeks following a fifth dose of mRNA vaccine, and analyzed by flow cytometry. The values represent the MFI normalized by CV3-25 mAb binding. (**C**) Neutralization activity was measured by incubating pseudoviruses bearing SARS-CoV-2 Spike glycoproteins with serial dilutions of plasma from vaccinated individuals. Neutralization half-maximal inhibitory serum dilution (ID_50_) values were determined using a normalized nonlinear regression using GraphPad Prism software. Limits of detection are plotted. (*** p<0.001; ****; p < 0.0001; ns: non-significant)

The Spike glycoprotein is a metastable fusion protein composed of a trimer of heterodimers expressed at the surface of viral particles which can be detected at the surface of infected cells (36-39). Its interaction with the receptor ACE2 on host cells enables S cleavage by host proteases, thus exposing the fusion peptide leading to viral entry (40-45). Given that the Spike glycoprotein is one of the main targets of humoral responses elicited by SARS-CoV-2 infection and vaccines, a strong selective immune pressure against this crucial protein led to the current evolution of emerging Omicron subvariants (21, 46-49). Each of these subvariants have acquired mutations in their Spike that help evade humoral responses, resulting in some cases in increased binding affinity for ACE2 (50-52). Interestingly, multiple Omicron lineages gained identical or similar Spike mutations in key antigenic sites in the receptor binding domain (RBD) and in the N-terminal domain (NTD), suggesting a convergent evolution (53, 54).

We previously demonstrated that temperature modulates the interaction between SARS-CoV-2 Spike and ACE2, with low temperature increasing ACE2 binding affinity and viral entry (55). We showed that this modulation was explained by favorable thermodynamic changes leading to the stabilization of the RBD-ACE2 interface and by the triggering a more ‘’open’’ conformation of the Spike trimer. Subsequent work on early Omicron subvariants (BA.1, BA.2, BA.2.12.1, BA.4/5) also showed an impact of temperature on Spike - ACE2 interaction (56). Here we extend these findings with recent Omicron subvariants which are characterized by enhanced immune escape and increased binding affinity to ACE2 compared to early Omicron strains. We tested the capacity of plasma from individuals who received a fifth dose of bivalent (BA.1 or BA.4/5) mRNA vaccine to recognize and neutralize several Spikes from recent Omicron subvariants. We next determined how temperature affects the interaction between Spike and ACE2 by combining an array of biochemical and biological assays, including biolayer interferometry, flow cytometry and virus capture assay. The associations between these parameters and the viral growth rate of each Omicron subvariants in the population between early 2022 and early 2024 was evaluated.

## MATERIALS AND METHODS

### Human Subjects

The study was conducted in 18 individuals, 11 females and 7 males (age range: 51–64 years). Blood samples were collected 22 days after the fifth dose of mRNA vaccine. We do not include other specific criteria, such as number of patients (sample size), sex, clinical, or demographic data, for admission to the cohort. Characteristics of the cohort are summarized in Table 1.

**Table 1.**
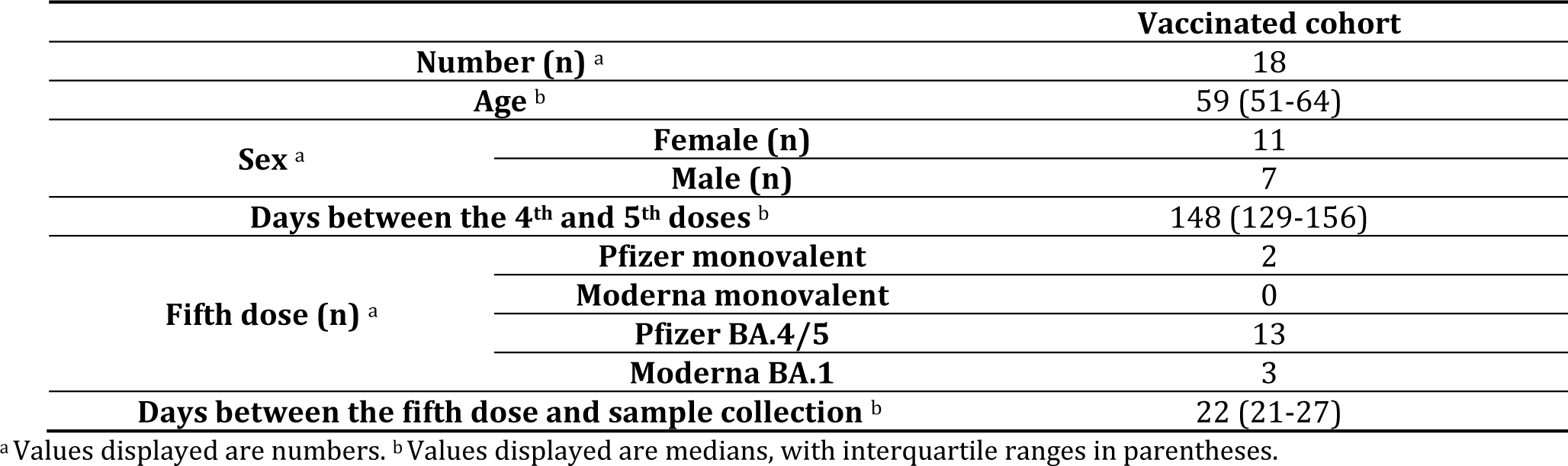
Characteristics of the SARS-CoV-2 vaccinated cohort.

|  |  | Vaccinated cohort |
| --- | --- | --- |
| Number (n) <sup>a</sup> |  | 18 |
| Age <sup>b</sup> |  | 59 (51-64) |
| Sex <sup>a</sup> | Female (n) | 11 |
|  | Male (n) | 7 |
| Days between the 4<sup>th</sup> and 5<sup>th</sup> doses <sup>b</sup> |  | 148 (129-156) |
| Fifth dose (n) <sup>a</sup> | Pfizer monovalent | 2 |
|  | Moderna monovalent | 0 |
|  | Pfizer BA.4/5 | 13 |
|  | Moderna BA.1 | 3 |
| Days between the fifth dose and sample collection <sup>b</sup> |  | 22 (21-27) |

<sup>a</sup> Values displayed are numbers. <sup>b</sup> Values displayed are medians, with interquartile ranges in parentheses.

### Plasmids

The plasmids expressing SARS-CoV-2 Spike D614G and SARS-CoV-2 RBD_WT_ (residues 319–541) fused with a 6xHis-tag were previously described (57). The RBD sequence (encoding for residues 319-541) fused to a C-terminal 6xHistag was cloned into the pcDNA3.1(+) expression vector. The plasmids encoding the full-length Spike from BA.2.75, CH.1.1, DV.7.1, BA.4/5, BQ.1.1, XBB, XBB.1, XBB.1.16, XBB.1.5, FD.1.1, EG.5.1, HK.3, BA.2.86 and JN.1 variants were generated by overlapping PCR using a codon-optimized wild-type SARS-CoV-2 Spike gene that was synthesized (Biobasic, Markham, ON, Canada) and cloned in pCAGGS as a template (16, 56, 58-60). All constructs were validated by Sanger sequencing. The plasmid encoding for soluble human ACE2 (residues 1–615) fused with an 8xHisTag was reported elsewhere (37). The plasmid encoding for the ACE2-Fc chimeric protein, a protein composed of an ACE2 ectodomain (1–615) linked to an Fc segment of human IgG1 was previously reported (61). The lentiviral vector pNL4.3 R^-^E^−^ Luc was obtained from NIH AIDS Reagent Program. The vesicular stomatitis virus G (VSV-G)-encoding plasmid was previously described (62).

### Cell lines

HEK293T cells (obtained from American Type Culture Collection [ATCC]) were derived from 293 cells, into which the simian virus 40 T-antigen was inserted. HEK293T-ACE2 were previously described (46) and were maintained in presence of puromycin (2µg/mL). Cf2Th cells (ATCC) are canine thymocytes resistant to SARS-CoV-2 entry and were used as target cells in the virus capture assay. HEK293T cells and Cf2Th were maintained at 37°C under 5% CO2 in Dulbecco’s Modified Eagle’s Medium (DMEM) (Wisent, St. Bruno, QC, Canada), supplemented with 5% fetal bovine serum (FBS) (VWR, Radnor, PA, USA) and 100 U/mL penicillin/ streptomycin (Wisent).

### Protein expression and purification

Proteins were expressed and purified as described (55). A detailed description is provided in the supplemental material.

### Flow cytometry analysis of cell-surface staining

Cell-surface staining of Spike glycoproteins was assessed by flow cytometry, as described (55). A detailed description is provided in the supplemental material.

### Virus capture assay

The SARS-CoV-2 virus capture assay was previously reported (56). A detailed description is provided in the supplemental material.

### Virus neutralization assay

Pseudoviral neutralization was performed as described (46, 57). A detailed description is provided in the supplemental material section.

### Western blot

At 48 h post-transfection, Spike-expressing HEK293T cells washed in PBS and then lysed with 1% Triton X-100 TNE lysis buffer (25 mM Tris (pH 7.5), 150 mM NaCl, 5 mM EDTA) supplemented with protease inhibitors cocktail (ThermoFisher scientific). Cell lysates were then centrifuged to pellet cell debris and cleared supernatants were transferred to new tubes. Laemmli buffer (Bio-Rad) with 5% B-mercaptoethanol (Bio-Rad) was then added to the cell lysates and subsequently heated at 95°C for 5min. Cell lysates were then resolved by SDS-PAGE and transferred to nitrocellulose membranes. Membranes were blocked for 1h at room temperature with blocking buffer (3% skim milk powder dissolved in tris-buffered saline supplemented with 0.1% Tween-20 [TBS-T]) and then probed with CV3-25 mAb (1ug/mL) followed by HRP-conjugated anti-human IgG (1:3000, Invitrogen). HRP enzyme activity was determined after the addition of a 1:1 mix of Western Lightning oxidizing and luminol reagents (PerkinElmer Life Sciences, Waltham, MA, USA), quantification of bands was performed using the ImageLab software (Bio-Rad).

### Cold inactivation

Pseudoviruses harboring the different S glycoproteins were produced as described in the supplemental material section, and small aliquots of each pseudovirus-containing supernatant were incubated on ice for different lengths of time (0h, 6h, 24h, 48h and 72h). Following incubation on ice, pseudoviruses were added in quadruplicate on HEK293T-ACE2 target cells which were seeded at a density of 1 × 10^4^ cells/well in 96-well luminometer-compatible tissue culture plates (PerkinElmer, Waltham, MA, USA) 24h before infection. Infectivity was measured as described above, with infectivity being relative to 0h on ice. The inactivation half-maximal inhibitory time (t_1/2_) represents the time needed to inhibit 50% of the infection of HEK293T-ACE2 cells by pseudoviruses.

### Biolayer interferometry

Binding kinetics were performed using an Octet RED96e system, as described (55, 56). A detailed description is provided in the supplemental material.

### Molecular dynamics simulations

MD simulations of BA.2.86 Spike were performed using NAMD 2.14 (63) and the CHARMM force field (64) using models of the Spike trimeric ectodomain based on the BA.2 structure in PDB entry 7XIX (65). For BA.2.86, three independent replica systems were generated, fully glycosylated, and fully solvated with TIP3P water using pestifer 1.2.9 (pypi.org/pestifer). Tertiary structural changes were monitored by measuring the distance between the center of mass of each RBD and the trimer center of mass. For BA.2.86, statistics on this distance are aggregated over all protomers within a trimer and over all trimer replicas.

### Lineage Growth Rate Estimation

We computed lineage growth rate estimates from GISAID metadata (downloaded January 8, 2024), based on pangolin annotation (66). We exclusively considered sequences labeled as “Omicron,” originating from human hosts, and featuring fully specified deposition and collection dates. Following Dadonaite et al. (50), we required sequences to have a collection date within 150 days of deposition, we excluded sequences with collection dates falling outside of 3.5 times the interquartile range of the median, and only sequences from countries with more than 500 samples and from lineages with more than 200 samples were retained. For the estimation of clade growth rates for the remaining 1170 lineages, we used a multinomial logistic regression model to analyze global lineage frequency data, accessible at the following repository: https://github.com/MurrellGroup/MultinomialLogisticGrowth/. Notably, our dataset includes analysis of emerging variants, such as JN.1, which differs from Dadonaite et al.’s results (50). In a comparative assessment of the 969 lineages analyzed by both methodologies, we obtained a near-perfect correlation with the recomputed estimates (Pearson correlation r=0.9987639, p-value < 2.2e-16), validating our results. Reported growth estimate for BA.4/BA.5 represents an average value derived from these two distinct lineages.

### Statistical analysis

Statistical analyses were done using GraphPad Prism version 8.4.2 (GraphPad). Every dataset was tested for statistical normality and this information was used to apply the appropriate (parametric or nonparametric) statistical analysis. P-values < 0.05 were considered significant; significance values are indicated as * p < 0.05, ** p < 0.01, *** p < 0.001, **** p < 0.0001.

## RESULTS

### Spike characteristics of recent Omicron sublineages

The diversification of Omicron sublineages is mainly driven through accumulation of mutations in its Spike glycoprotein (19). Compared with BA.2, BA.2.75 possesses 9 additional mutations in its Spike (Figure 1A), conferring improved ACE2 binding and higher evasion from vaccine-elicited and monoclonal antibodies (mAbs). It was demonstrated that two mutations in the RBD, G446S and N460K, had a profound effect (23, 67-69). CH.1.1, one of the most frequently observed subvariant derived from BA.2.75, gained more attention in Asia and Europe (Figure 2A) (22). Compared to BA.2.75, CH.1.1 has four additional mutations in its RBD, R346T, K444T, L452R and F486S (Figure 1A). These mutations have been reported to increase neutralization escape from polyclonal serum and mAbs (16, 22, 70, 71). Interestingly, L452R, which is also present in the Spike of BA.4, BA.5 and their progeny BQ.1.1, also improves RBD and Spike affinity to the ACE2 receptor (52, 72-74). BQ.1.1 harbors three additional antibody-escape mutations in its RBD compared to BA.4/BA.5: R346T, K444T and N460K (14, 24, 53, 75, 76). While infections involving XBB were increasing due to its improved antibody evasion (16, 25, 76-78), XBB.1.5 quickly dominated over XBB and BQ.1.1, due to its improved ACE2 affinity over XBB (27-29) and improved immune escape compared to BQ.1.1 (79-81).

**Figure 2.**
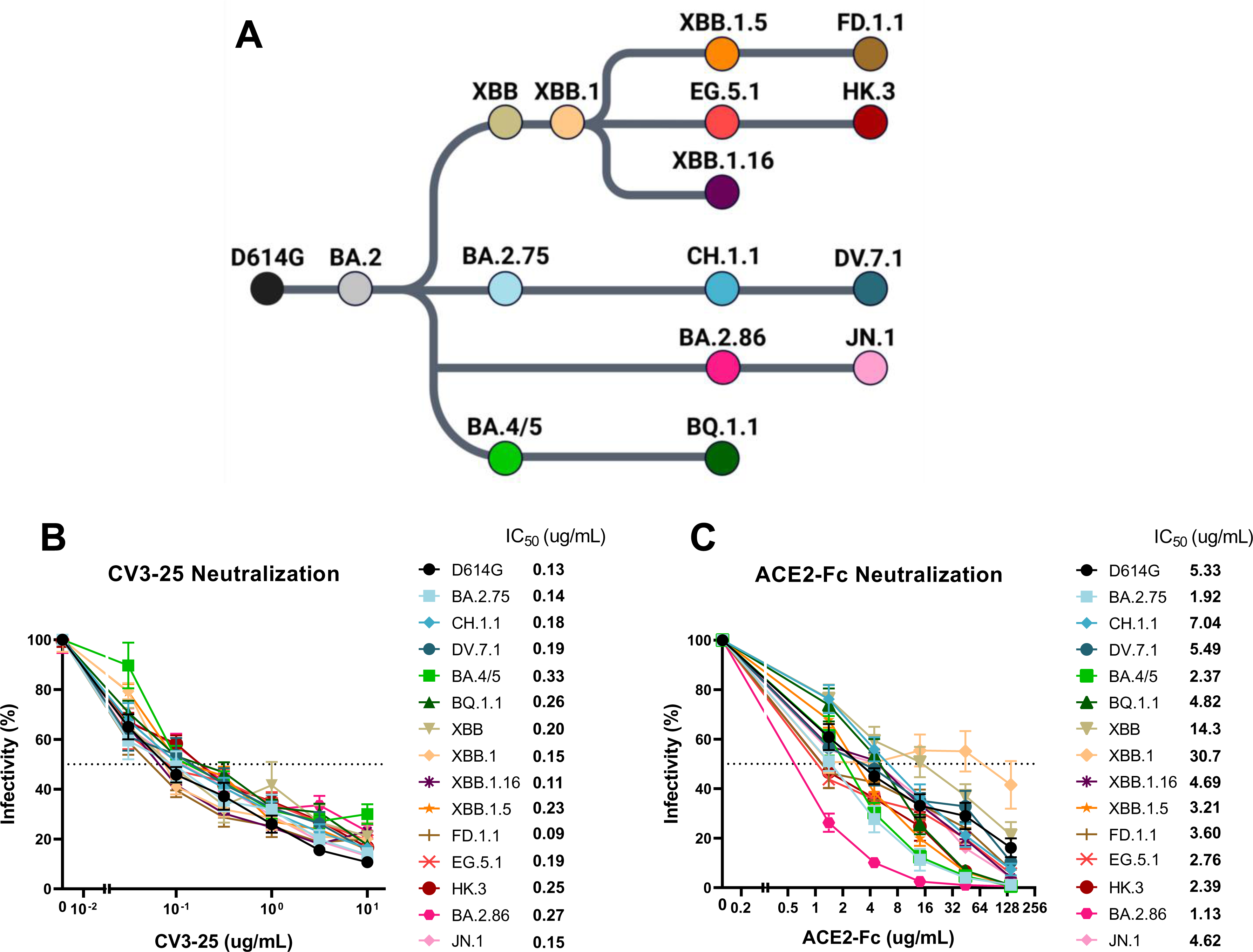
Evolution of Omicron subvariants and their susceptibility to neutralization by CV3-25 mAb and ACE2-Fc. (**A**) Schematics showing the evolving Omicron sublineages of predominant strains. (**B and C**) Neutralization activity was measured by incubating pseudoviruses bearing SARS-CoV-2 Spike glycoproteins with serial dilutions of anti-S2 CV3-25 monoclonal antibody (**B**) or ACE2-Fc (**C**). Neutralization half-maximal protein concentration (IC_50_) values were determined using a normalized nonlinear regression using GraphPad Prism software.

SARS-CoV-2 descendants of XBB.1 with the S:S486P mutation (Figure 1A, 2A), also known to enhance ACE2 affinity (27), are predominantly circulating worldwide (30). Notably, XBB.1.16, harboring in its Spike the E180V, S486P and T478R substitutions, showed a remarkable growth advantage compared to XBB.1.5 although both Spike glycoproteins possess similar characteristics in terms of infectivity and immune escape (82, 83). Predominant lineages possess a substitution at either or both positions S:L455F and S:F456L, mostly known as the ‘’FLip’’ mutations (L455F+F456L) (31, 84, 85). The ‘’FLip’’ mutations have been shown to act synergistically to further enhance ACE2 binding affinity and neutralization escape from plasma (84). Compared to XBB.1.5, FD.1.1 harbors the F456L substitution, EG.5.1 has the Q52H and F456L substitutions while its sublineage named HK.3 also acquired the L455F mutation. DV.7.1, a sublineage of CH.1.1, also started spreading which may be due to the acquisition of the ‘’FLip’’ mutations (Figure 1A) (84, 86). BA.2.86, a highly divergent BA.2 subvariant, possesses mutations in its Spike enhancing its immune evasion and ACE2 binding compared with XBB.1.5, notably N450D, K356T, L452W, A484K, V483del and V445H, with the addition of R403K being responsible for its enhanced ACE2 binding affinity (33, 87-90) (Figure 1A, 2A). Lastly, the BA.2.86 subvariant, JN.1, possesses an additional substitution at S:L455S, conferring improved immune evasion in spite of reduced ACE2 binding (30, 32, 35).

### Plasma recognition and neutralizing activity

To monitor the humoral responses against recent Omicron subvariants, we collected plasma samples from 18 donors (11 females and 7 males), with a total median age of 59 years (interquartile range: 51-64 years) (Table 1). Plasma samples were collected three to four weeks after the fifth dose of mRNA vaccine with 13/18 receiving the Pfizer mRNA BA.4/5 bivalent vaccine, 2/18 receiving the Pfizer monovalent vaccine and 3 receiving the Moderna mRNA BA.1 bivalent vaccine (Table 1).

We first evaluated the capacity of plasma to recognize recent Omicron subvariants by transfecting HEK293T cells with plasmids encoding the full-length SARS-CoV-2 Spikes (52, 91). The CV3-25 monoclonal antibody (mAb) was used for the normalization of Spike expression levels across experimental conditions. CV3-25 is a Spike-binding mAb, specific against a conserved epitope shared among β-coronaviruses within the S2 subunit. (58, 92-94). We observed that all tested Omicron subvariant Spikes were less efficiently recognized by plasma than the D614G Spike, with DV.7.1 (x3.5), HK.3 (x2.8) and JN.1 (x2.7) Spikes demonstrating the highest fold change in recognition by plasma (Figure 1B). Interestingly, BA.2.86 (x2.0) was better recognized than CH.1.1 (x2.5), XBB.1.5 (x2.2) and EG.5.1 (x2.2). As expected, BA.4/5 (x1.6), BQ.1.1 (x1.8) and BA.2.75 (x1.9) had a decreased ability to evade recognition by plasma from vaccinated individuals compared to BA.2.75- and XBB-sublineages (Figure 1B).

We next measured the capacity of plasma to neutralize pseudoviral particles harboring Spikes from emerging Omicron sublineages. To do this, we produced pseudoviral particles bearing the Spike glycoprotein from several omicron subvariants, as previously described (46, 47, 95). As expected, all plasma strongly neutralized pseudoviral particles bearing the D614G Spike after the fifth dose of mRNA vaccine (Figure 1C). In agreement with Spike recognition by plasma (Figure 1B), all Omicron subvariants demonstrated significant neutralization escape, with the DV.7.1 (x66.8) and HK.3 (x57.1) Spikes being the most resistant to neutralization and the BA.2.75 (x1.4) and BA.4/5 (x.1.4) Spikes the most efficiently neutralized by plasma from vaccinated individuals. Interestingly, we observed a slight enhanced neutralization escape with BA.2.86 (x12.2) compared to most XBB sublineages, which was further pronounced with JN.1 (x18.0) (Figure 1C).

### Neutralization susceptibility to CV3-25 and ACE2-Fc

As a control for our neutralization experiments, we first measured the susceptibility of the different variants to the S2-targeting CV3-25 mAb which recognizes a highly conserved epitope (58, 92-94). As expected, all subvariants tested were similarly susceptible to CV3-25 neutralization, with an IC_50_ in the range of 0.09-0.33 µg/mL (Figure 2B). Given that recent Omicron subvariants have acquired mutations known to enhance ACE2 affinity, we next evaluated the ability of recombinant human ACE2-Fc to neutralize these subvariants using pseudoviral particles (61). We found that most subvariants had improved susceptibility to ACE2-Fc neutralization compared to D614G (Figure 2C). However, XBB and XBB.1 were remarkedly less susceptible to ACE2-Fc neutralization compared to D614G. Interestingly, CH.1.1 and DV.7.1 were less sensitive to ACE2-Fc compared to their parental strain BA.2.75, although an improvement was seen for DV.7.1, possibly owing to the ‘’FLip’’ mutations (Figure 1A, 2C). As expected, XBB sublineages harboring the S:S486P were neutralized more efficiently than the parental XBB.1 strain, with EG.5.1 and HK.3 being the most susceptible (Figure 1A, 2A, 2C). The additional mutations in BQ.1.1 decreased its susceptibility to ACE2-Fc-mediated neutralization compared to BA.4/5. Of note, among all the Omicron subvariants tested, BA.2.86 was the most susceptible to ACE2-Fc neutralization, with a 4.7x fold decrease in IC_50_ compared to D614G (Figure 2C). The additional substitution in JN.1 considerably reduced its sensitivity to ACE2-Fc neutralization compared to BA.2.86.

### Impact of temperature on the RBD-ACE2 interaction

Previous studies showed that SARS-CoV-2 Spike interaction with ACE2 can be modulated by temperature, with lower temperatures enhancing RBD affinity for ACE2 (55, 56). To evaluate whether this property was conserved with these new Omicron subvariants, we measured the RBD affinity to soluble ACE2 (sACE2) by performing biolayer interferometry (BLI) experiments at different temperatures (35°C, 25°C and 15°C). In agreement with previous observations (52, 55), we found that cold temperatures enhanced the affinity of the RBDs tested with ACE2 (Figure 3A). The higher affinity observed at colder temperatures for the different Omicron subvariants RBDs compared to wild-type (WT) RBD was mainly explained by a drastic decrease in the off rate (Figure S1A). Interestingly, except for RBD_XBB_, the RBD of Omicron subvariants had similar or better affinity for ACE2 at 25°C compared to RBD_WT_ at 15°C, and these differences were further enhanced for all Omicron subvariants at 15°C. The RBD_BA.4/5_ and RBD_BQ.1.1_ affinity was relatively similar at all temperature tested. In agreement with our ACE2-Fc neutralization results, RBD_BA.2.86_ had the highest affinity for sACE2 among the different Omicron subvariants and this remained true for all temperature tested.

**Figure 3.**
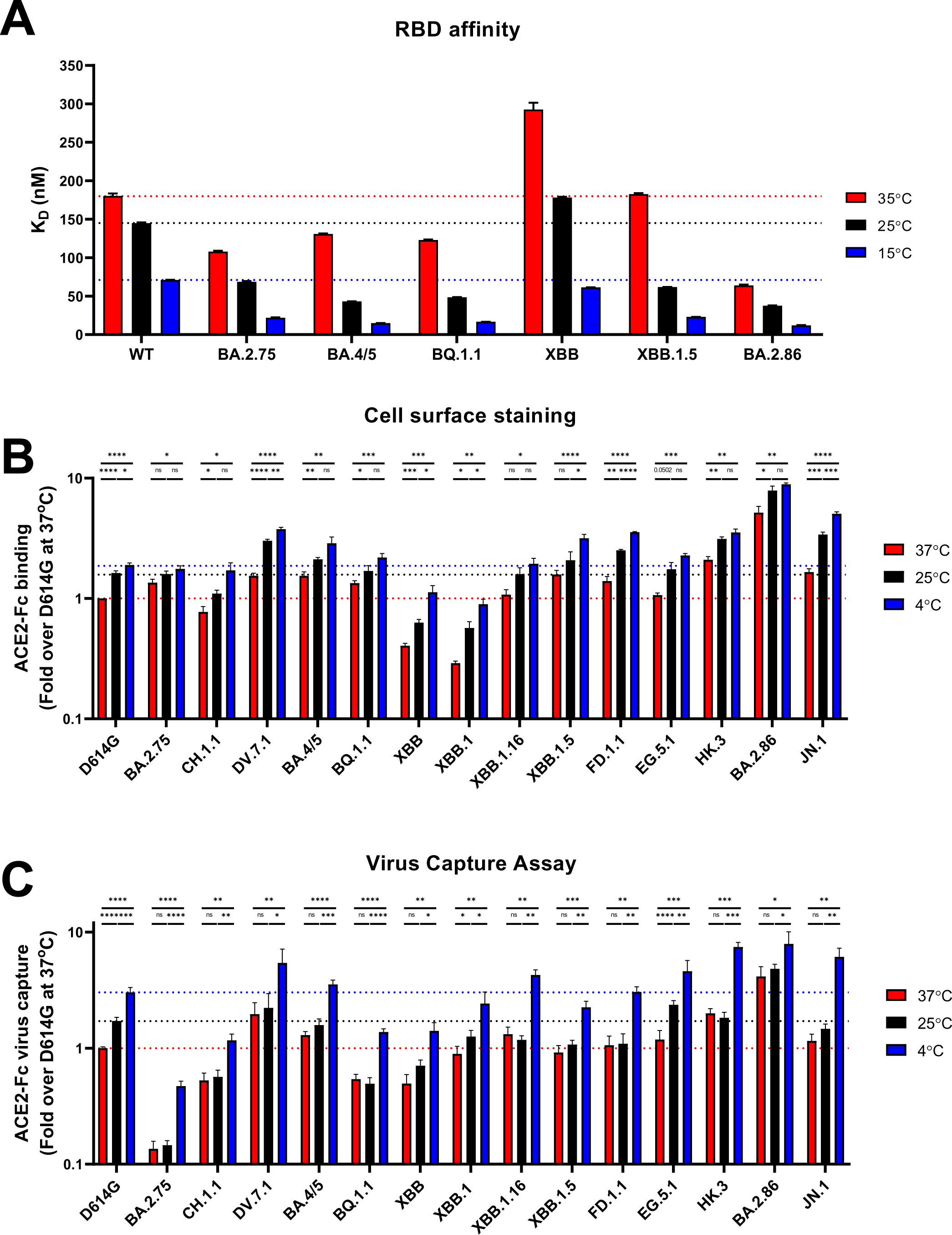
Low temperature enhances Omicron subvariants RBD affinity and Spike interaction with ACE2. (**A**) Binding kinetics between RBD_WT_ and several RBD_Omicron_ with sACE2 assessed by BLI at different temperatures. Biosensors loaded with RBD proteins were soaked in twofold dilution series of sACE2 (500 nM–31.25 nM) at different temperatures (35°C, 25°C or 15°C) with the affinity constants (K_D_) being represented along with the technical error. Affinity values obtained at different temperatures were calculated using a 1:1 binding model. (**B and C**) Cell surface staining of Spike-expressing HEK293T cells and virus capture assay of WT (D614G) and Omicron subvariant Spikes at 37°C (red), 25°C (black), and 4°C (blue). (**B**) ACE2-Fc recognition is presented as a ratio of ACE2 binding over D614G Spike obtained at 37°C. (**C**) Pseudoviruses encoding the luciferase gene (Luc+) and expressing the several SARS-CoV-2 Spikes were tested for viral capture by ACE2-Fc at the respective temperatures. Relative light units (RLU) obtained using ACE2-Fc was normalized to the signal obtained with the temperature-independent CV3-25 mAb and presented as a ratio of ACE2 capture to D614G obtained at 37°C. (**B and C**) These results represent at least three independent experiments showing means ± SEM. Statistical significance was tested using unpaired T tests (**B and C**). (* p < 0.05; ** p < 0.01; *** p < 0.001, **** p < 0.0001, ns: non-significant)

Moreover, the RBD of BA.2.75, BA.4/5, BQ.1.1 and BA.2.86 bound better to ACE2 compared to RBD_WT_ at the three temperatures tested (Figure 3A). Of note, the affinity of RBD_XBB_ was remarkedly lower compared to RBD_WT_, but this was restored with the acquisition of S:S486P as shown with RBD_XBB.1.5_, thus corroborating the ACE2-Fc neutralization results observed for XBB/XBB.1 and XBB.1.5 (Figure 2C).

### Impact of temperature on Spike-ACE2 interaction

We next measured the capacity of ACE2-Fc to bind the full-length Omicron subvariants Spikes expressed at the surface of transfected HEK293T cells using a well-established flow cytometry assay (52, 55, 56, 61) (Figure 3B). We used the temperature-independent S2-targeting CV3-25 mAb as a control for each temperature (55, 56, 92) (Figure S1B). Lower temperatures have been shown to modulate the capacity of the trimeric Spike to interact with ACE2 by favoring the adoption of the RBD ‘’up’’ conformation, required for ACE2 binding (45, 55, 56). To extend these results on recent Omicron subvariants, we evaluated ACE2-Fc binding with Spike-expressing cells after incubation at 37°C, 25°C and 4°C (Figure 3B). In agreement with the results obtained by BLI (Figure 3A, S1C), we observed a gradual increase in ACE2-Fc binding concomitant with the temperature decrease for all Spikes tested. Of note, DV.7.1, BA.4/5, XBB.1.5, FD.1.1, HK.3,BA.2.86 and JN.1 showed better binding at 25°C than D614G at 4°C, and these differences were further enhanced at 4°C. Interestingly, HK.3 and BA.2.86 showed enhanced binding at 37°C compared to D614G binding at 4°C. As expected from previous reports showing higher affinity for RBD interaction (32, 33), BA.2.86 presented the highest binding capacity at all temperatures tested, showing more than 8.8-fold improvement at 4°C compared to D614G at 37°C. As expected, JN.1 showed reduced ACE2-Fc binding compared to its parental lineage BA.2.86. In addition, DV.7.1, HK.3 and BA.2.86 had the highest ACE2-Fc binding capacity at 37°C, with BA.2.86 demonstrating the greatest binding (Figure 3B). The acquisition of S:S486P improved ACE2-Fc binding for XBB.1.16, XBB.1.5, FD.1.1 and EG.5.1 compared to XBB/XBB.1, independently of the temperature.

We next evaluated the impact of temperature on ACE2 interaction with Spike expressed at the surface of pseudoviral particles. To investigate this, we used a previously described virus capture assay (55, 56, 96). Briefly, we produced pseudoviral particles bearing the different Omicron subvariant Spikes and evaluated their ability to interact with ACE2-Fc on ELISA plates. Pseudoviral particles were pre-incubated at different temperatures (37°C, 25°C and 4°C). In agreement with enhanced RBD affinity and Spike interaction with ACE2 at lower temperatures (Figure 3A-B), we observed a stepwise increase in viral capture at colder temperatures for all Spikes tested, although the increase at 25°C was modest for most Spikes (Figure 3C). These results significantly correlated with the RBD affinity and cell-based binding assay at different temperatures (Figure S1C, S1D). Interestingly, we also saw an improved binding for DV.7.1, HK.3 and BA.2.86 at 37°C, which was higher than that of D614G at 25°C, with BA.2.86 showing the greatest capture by ACE2 at all respective temperatures tested. On the other hand, JN.1 showed a marked capture at 4°C, despite having reduced virus capture at 37°C and 25°C. Altogether, our findings demonstrate that Spike-ACE2 interaction of recent Omicron subvariant is modulated by temperature independently of whether the Spike is expressed on pseudoviral particles or at the cell surface, likely explained through higher RBD affinity.

Of note, at 37°C most Omicron subvariants bound ACE2-Fc better than D614G (Figure 3C). We also noticed a decrease in ACE2 capture for BQ.1.1 and JN.1 compared to BA.4/5 and BA.2.86 respectively, which corroborated the results obtained by ACE2-Fc neutralization and binding (Figure 2B, 3B). Interestingly, XBB.1 but not XBB had a similar capture by ACE2-Fc compared to D614G. As expected, lineages harboring the S:S486P substitution had improved capture by ACE2-Fc, and this was more noticeable for XBB.1.16 (Figure 3C). Finally, DV.7.1, HK.3 and BA.2.86 had the highest binding at 37°C to immobilized ACE2, with BA.2.86 showing the greatest binding among all the Omicron subvariants tested (Figure 3C).

### Impact of temperature on ACE2 binding cooperativity

Since Spike interaction with ACE2 requires RBD to be in the ‘’up’’ conformation (39, 97, 98), we wondered whether recent Omicron subvariants had an improved propensity to adopt the ‘’up’’ conformation at lower temperature as previously observed for SARS-CoV-2 D614G and early omicron strains (55, 56, 99). To investigate this, we calculated the Hill coefficient (h), which is a measure indicating the degree of binding cooperativity between protomers of a multimeric protein with its ligand (100, 101). Briefly, we transfected HEK293T cells with the full-length Spikes of recent Omicron subvariants and incubated these cells with increasing concentrations of monomeric soluble ACE2 as described previously (55, 56). The Hill coefficient was calculated based on the steepness of the doses-responses curves, as described in the material and methods section.

Consistent with previous observations, the Hill coefficient at low temperature (4°C) was greater than at 37°C for all Spikes tested, confirming that low temperatures facilitate ACE2- induced Spike opening (Figure 4) (55, 56). Compared to D614G, all Omicron subvariants showed enhanced cooperativity at 4°C, with a striking improvement for HK.3, BA.2.86 and JN.1 (Figure 4M-O). Interestingly, CH.1.1 and DV.7.1 showed similar cooperativity at 4°C. Consistent with their improved binding and cooperativity at 37°C, XBB lineages harboring the S:S486P had enhanced cooperativity at 4°C compared to XBB/XBB.1, with XBB.1.5 having the greatest improvement (Figure 4G-L).

**Figure 4.**
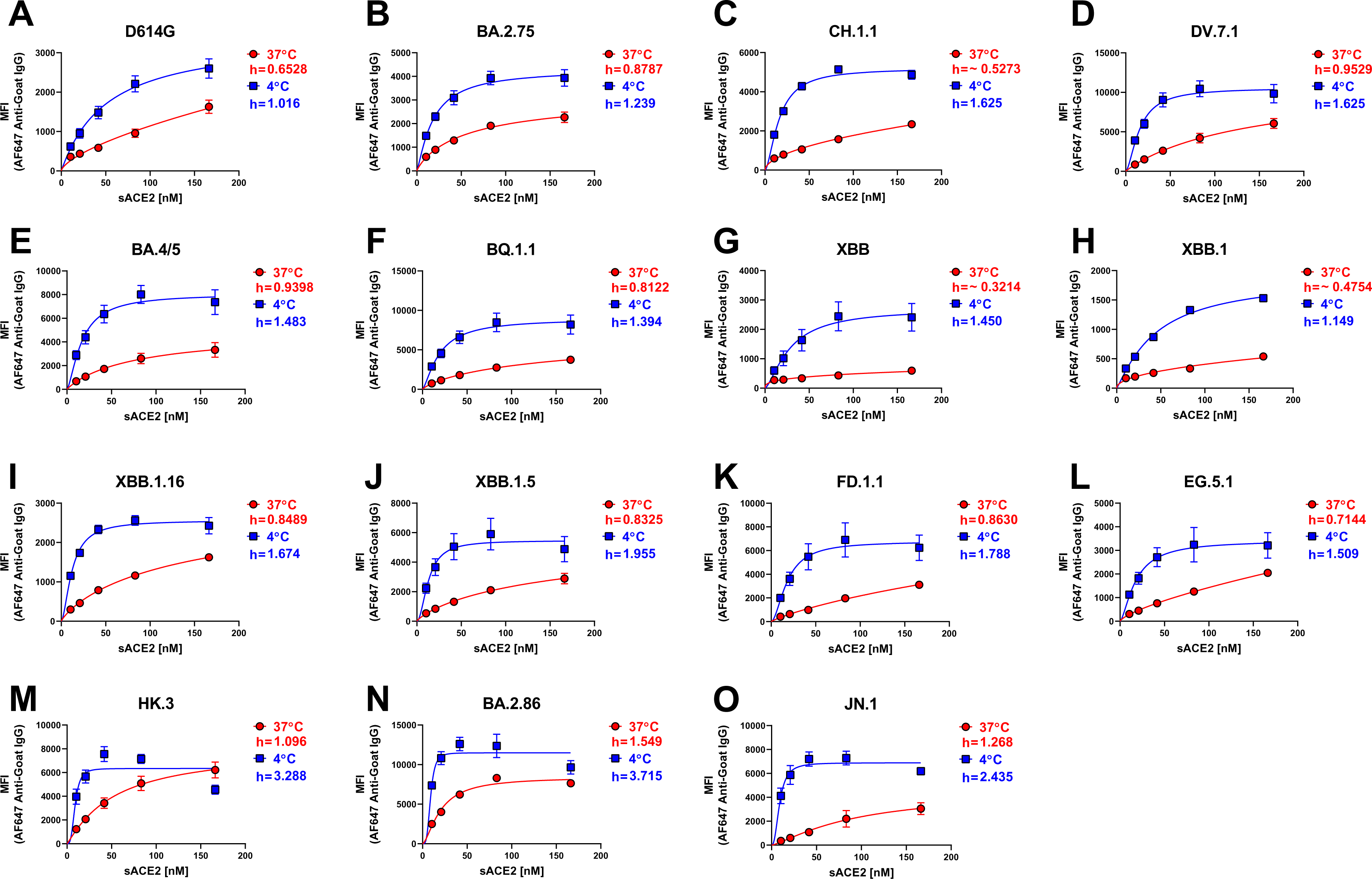
Omicron subvariant Spikes “open-up” at low temperature. Binding of sACE2 to the Spike of D614G and several Omicron subvariants expressed at the surface of HEK293T cells was measured by flow cytometry. Increasing concentrations of sACE2 were incubated with Spike-expressing cells at 37°C (red) or 4°C (blue). Means ± SEM derived from at least three independent experiments are shown. Hill coefficients were calculated using the GraphPad software.

We also observed that at 37°C, most Omicron subvariants had a higher Hill coefficient than D614G (Figure 4), except for CH.1.1, XBB and XBB.1 (Figure 4A, 4C, 4G-H). Consistent with the ACE2-Fc binding at the cell surface and viral particles (Figure 3B-C), XBB.1.16, XBB.1.5, FD.1.1 and EG.5.1 demonstrated a marked improvement in promoter cooperativity compared to XBB.1, likely due to the S:S486P substitution (Figure 4G-L). Aligned with their improved ACE2- Fc binding, DV.7.1, HK.3 and BA.2.86 showed the best cooperativity at 37°C among all the Omicron subvariants tested (Figure 4D, 4M-N). Interestingly, despite lower ACE2-Fc binding at the surface of transfected cells and pseudoviruses, JN.1 shows high levels of cooperativity at 37°C (Figure 4O). These results suggest that Omicron subvariants acquired mutations improving their Spike opening.

### Spike processing is associated with susceptibility to cold inactivation

It has been demonstrated that Omicron BA.1 Spike processing is reduced compared to D614G and Delta, and that concomitantly, BA.1 Spike is more susceptible to cold inactivation (102). To test the hypothesis that Spike cleavage could be associated with susceptibility to cold inactivation, we first measured Spike processing at the cell surface (Figure 5A). HEK293T cells were transfected with the Omicron subvariants Spikes and subsequently examined by western blotting using the anti-S2 CV3-25 mAb (Figure 5A). Compared with D614G, most Omicron subvariants exhibited similar or higher cleavage of S to S2 (S2/S ratio), with the exception of BA.2.75 whose processing was decreased. Interestingly, CH.1.1 and BQ.1.1 had an enhanced processing compared to their respective parental lineage. Among the XBB.1 sublineages, XBB.1.5 and EG.5.1 exhibited the highest level of processing, with HK.3 showing reduced processing. BA.2.86 and JN.1 presented improved processing compared to D614G and XBB.1 sublineages.

**Figure 5.**
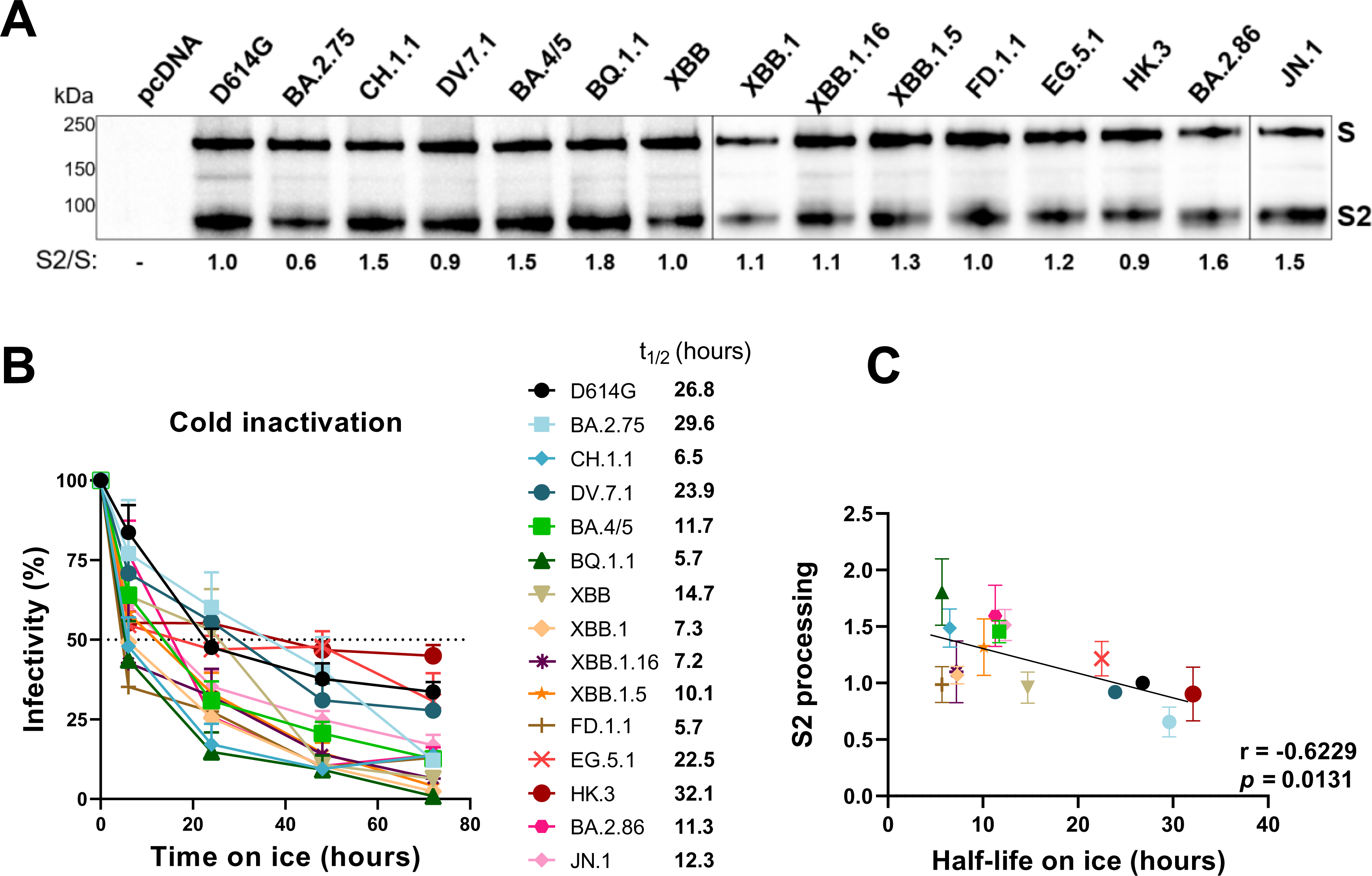
Omicron subvariants’ Spike processing and susceptibility to cold inactivation. (**A**) HEK293T cells were transfected with several Omicron subvariants Spikes, which were subsequently lysed and analyzed by western blot using the anti-S2 CV3-25 monoclonal antibody. The ratio of processed S2 over unprocessed S are shown (S2/S), with values being normalized to D614G for each experiment. Values represents the mean of at least three independent experiments. (**B**) To evaluate the susceptibility of Omicron subvariants to cold inactivation, pseudoviral particles bearing the several Spikes were incubated on ice for 0h, 6h, 24h, 48h and 72h, followed by infection of HEK293T-ACE2 cells. These results represent at least three independent experiments showing means ± SEM. The time required to reduce infection by 50% (t_1/2_) was determined using a normalized nonlinear regression using GraphPad Prism software. (**C**) Pearson rank correlations between the S2/S ratio with the half-life on ice.

To investigate the role of S2 processing in the susceptibility of recent Omicron subvariants to cold inactivation, we next evaluated the effect of cold (0°C) on the infectivity of pseudoviral particles (102-105). Briefly, pseudoviral particles bearing the different Omicron subvariants Spikes were incubated on ice for various amounts of time and their capacity to infect HEK293T- ACE2 cells was subsequently measured (Figure 5B). Interestingly, except for BA.2.75 and HK.3, all Omicron subvariants were more susceptible to cold inactivation compared to D614G, with FD.1.1 and BQ.1.1 being the most susceptible. Compared to CH.1.1 and EG.5.1, the acquisition of the ‘’FLip’’ mutations seemed to improve the stability of DV.7.1 and HK.3 Spikes respectively. We found a negative correlation between S2 processing (S2/S) and the half-life on ice (t_1/2_), suggesting that enhanced processing is associated with higher susceptibility to cold inactivation (Figure 5C).

### MD simulations of BA.2.86 Spikes are consistent with more open trimers at lower temperatures

To corroborate our findings that lower temperatures improve the propensity of Omicron subvariants RBD to adopt the ‘’up’’ conformation, all-atom, explicit-solvent MD simulations were performed on three independent replicas of BA.2.86 homology models based on the closed BA.2 spike structure at both 4°C and 37°C (Figure 6). The degree of opening of the RBDs was assessed by measuring the instantaneous distance between RBD center of mass and the center of mass of the full trimer, as described (55). Plots of this distance vs simulation time and their empirical distributions are shown in Figure 6B. The BA.2.86 systems required more than 150 ns of equilibration, after which the colder spikes showed distances of about 48 Å and warmer spikes about 47 Å. The Wuhan strain spike considered in our earlier work (55) showed distances of about 46 Å at 4°C and 45 Å at 37°C, which is a similar sensitivity to temperature but with overall more closed-down RBDs relative to BA.2.86. The equilibrium empirical distributions of these distances show that the warmer spikes sample three distinct states while the colder spikes are more uniformly in a single, more open state (Figure 6).

**Figure 6.**
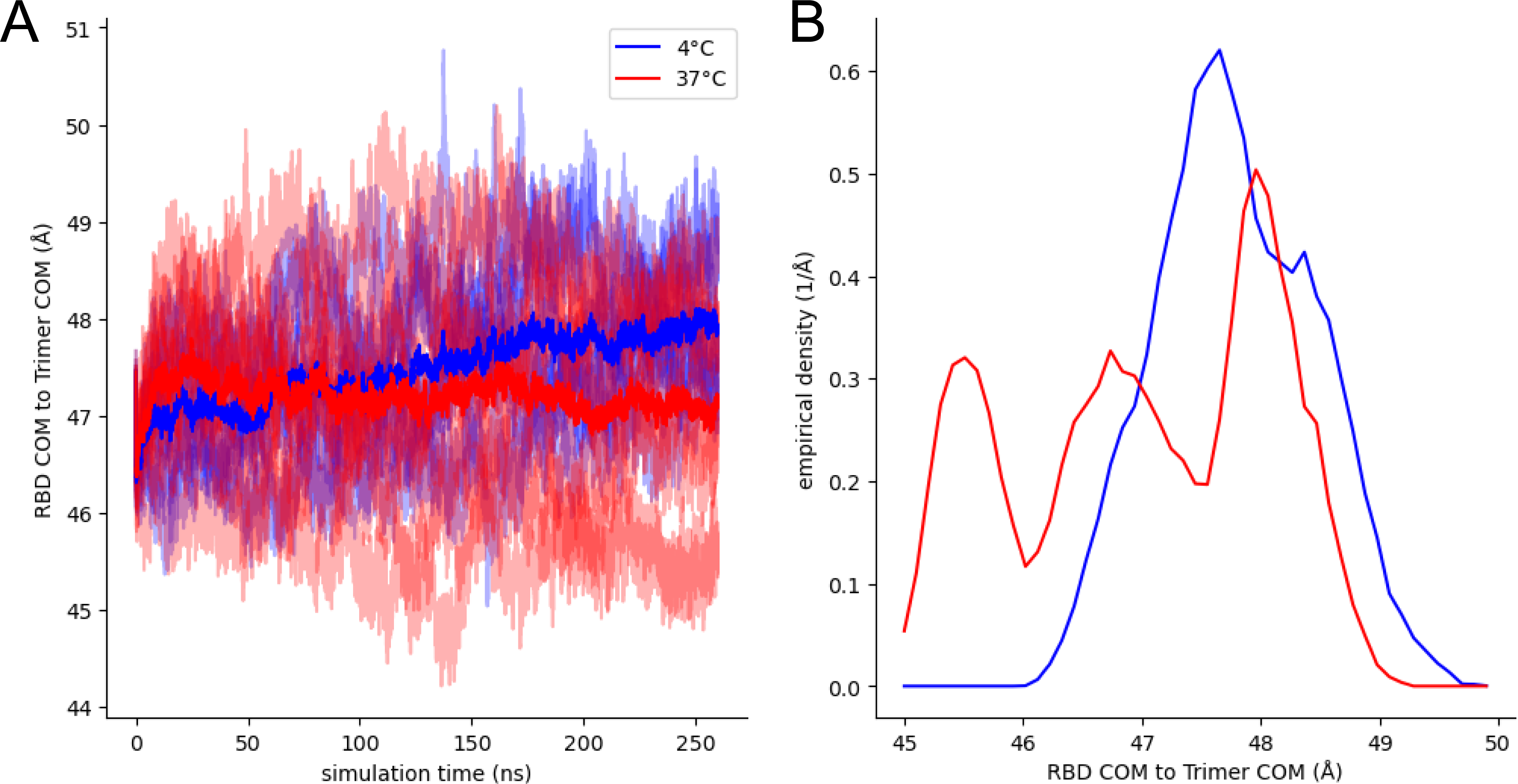
MD simulations of BA.2.86 Spike are consistent with more open trimers at lower temperatures. (**A**) Traces of the distances between RBD and trimer centers of mass (COM) from three replicas each of all-atom, fully glycosylated, and solvated MD simulations of the closed, BA.2.86 SARS-CoV-2 S trimers at 4°C (blues) and 37°C (reds) with dataset averages shown in heavy traces. Each set has nine distinct traces. (**B**) Empirical density distributions of these distances sampled after 150 ns of simulation time.

### ACE2 interaction at the surface of viral particles is associated with viral growth rates

SARS-CoV-2 viral transmission is a complex multifactorial process where immune pressure, tropism and affinity for its receptor affect the outcome (5). We investigated whether Spike recognition, either at the surface of cells or viral particles, as well as ACE2 interaction was associated with viral growth rates. To do this, we calculated the growth rates of SARS-CoV-2 Omicron sublineages following the methodology developed by Dadonaite et al. (50) to incorporate recently emerged Omicron subvariants as of January 8^th^ 2024. We found that plasma recognition of Omicron subvariants Spikes expressed at the cell surface was significantly associated with the growth rate of the different Omicron subvariants (Figure 7A); but not binding to ACE2 at 37°C (Figure 7B). Interestingly, plasma neutralization of pseudoviral particles bearing emerging Omicron subvariants Spikes was significantly associated with viral growth rates (Figure 7E). Intriguingly, the capacity of the different Spikes to interact with ACE2, particularly at low temperature was significantly associated with the growth rates (Figure 7C-D, F-H). Higher associations were observed when the Spike was expressed at the surface of viral particles rather than at the cell surface (Figure 7H vs 7D). Of note, the combination of both recognition escape or neutralization escape with ACE2-Fc binding at low temperatures was better associated with viral growth rates (Figure 7I,J). Thus, suggesting that antibody escape and ACE2 interaction at low temperatures contribute to viral transmission.

**Figure 7.**
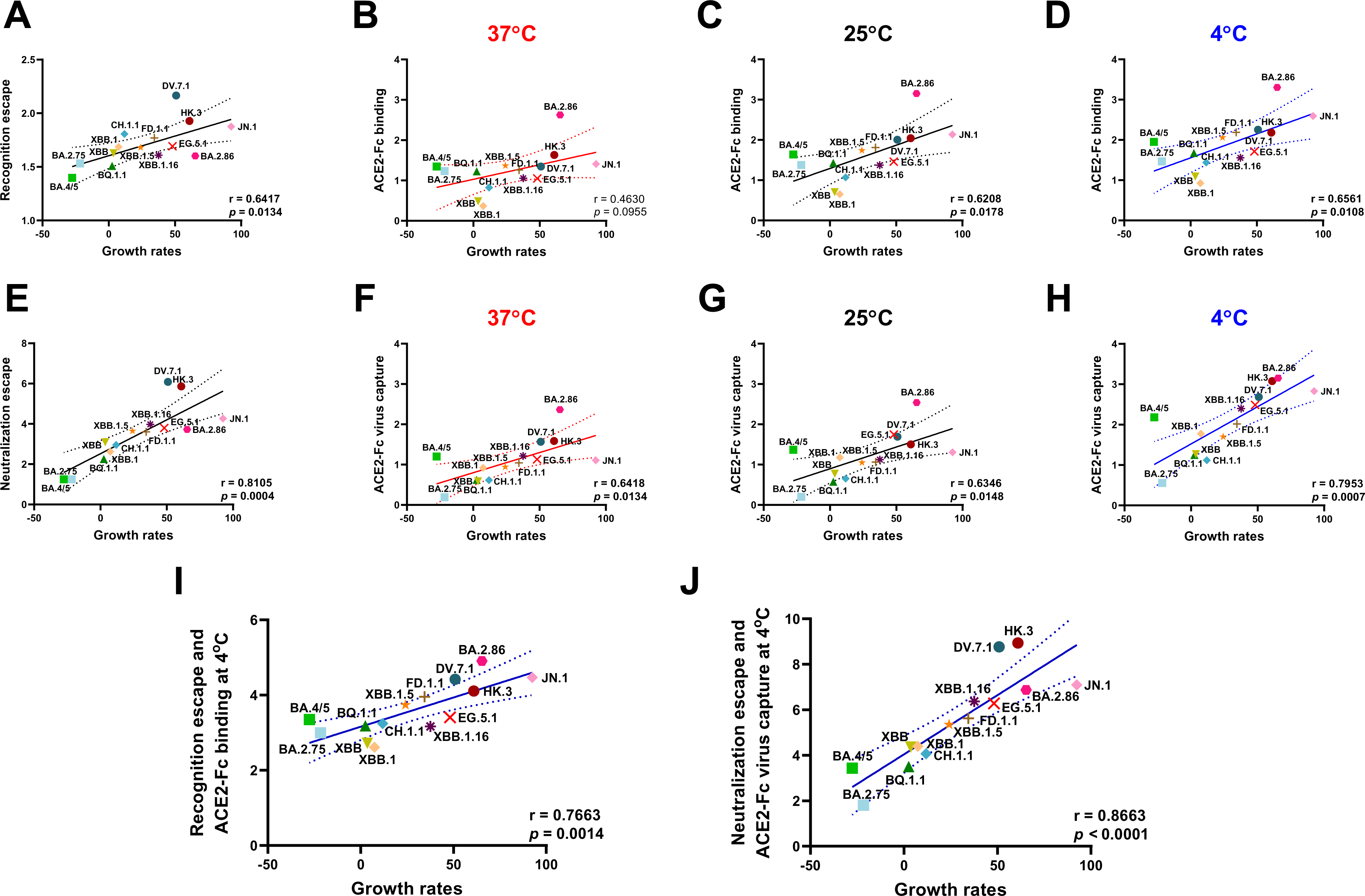
Association of immune escape and ACE2 interaction with Omicron subvariants growth rates. Associations between growth rates of several Omicron subvariants and immune escape or ACE2 interaction at different temperatures. Correlation of growth rates with plasma recognition (**A**) or ACE2-Fc binding to Spikes presented at the cell surface at 37°C (**B**), 25°C (**C**) and 4°C (**D**). Correlation of growth rates with plasma neutralization (**E**) or ACE2-Fc virus capture at 37°C (**F**), 25°C (**G**) and 4°C (**H**). Plasma recognition and neutralization escape, ACE2-Fc binding and ACE2-Fc virus capture were normalized to the values obtained for D614G at 37°C and subsequently log_2_ transformed. Panel I and J represents correlations using the combination of plasma recognition or neutralization escape with ACE2-Fc binding at the surface of cell or viral particles. (**A - J**) Pearson rank correlations are shown. Panels A, E, I and J refers to data shown in Figure 1. Panel B-D, F-H and I-J refers to data shown in Figure 3. (* p < 0.05; ** p < 0.01; *** p < 0.001, **** p < 0.0001)

## DISCUSSION

The continued evolution of SARS-CoV-2 requires constant monitoring of its new variants (21). Current circulating strains are derived from the Omicron variant, with each newly emerging subvariant showing improved transmission which can mostly be explained by their enhanced antibody escape and ACE2 binding affinity (54, 106). Here we characterize the functional properties of recent Omicron subvariant Spikes, including their capacity to evade humoral responses. The phenotype of the various Omicron subvariants Spikes are summarized in Table 2.

**Table 2.** Omicron subvariants Spikes characteristics. Summarized data shown in Figures 1, 2, 3, 4, 5 and 7 were plotted in a table. Relative to D614G, a decrease in the parameter measured are shaded orange to a degree dependent on the level of the decrease. Increases relative to D614G for the parameters measured are shaded green to a degree dependent on the level of the increase.

| SARS-CoV-2 S glycoprotein | Growth rates<br>This study | Plasma post 5th dose |  | Neutralization (IC<sub>50</sub>) |  | RBD affinity (nM) |  |  | Relative ACE2-Fc binding |  |  | Relative ACE2-Fc Virus capture |  |  | Hill coefficient |  | Western blot | Cold inactivation |
| --- | --- | --- | --- | --- | --- | --- | --- | --- | --- | --- | --- | --- | --- | --- | --- | --- | --- | --- |
|  |  | Plasma binding (Normalized to CV3-25) | Neutralization (ID<sub>50</sub>) | ACE2-Fc (ug/mL) | CV3-25 (ug/mL) | 35°C | 25°C | 15°C | 37°C | 25°C | 4°C | 37°C | 25°C | 4°C | 37°C | 4°C | Relative S2 processing | t<sub>1/2</sub> (hours) |
| D614G | n/a | 99.2 | 5480.7 | 5.33 | 0.13 | 180.20 | 144.90 | 71.08 | 1.00 | 1.59 | 1.88 | 1.00 | 1.72 | 3.03 | 0.65 | 1.02 | 1.0 | 26.8 |
| BA.2.75 | -21.6 | 52.4 | 3971.3 | 1.92 | 0.14 | 107.90 | 68.46 | 21.80 | 1.35 | 1.60 | 1.76 | 0.14 | 0.15 | 0.47 | 0.88 | 1.24 | 0.7 | 29.6 |
| CH.1.1 | 11.8 | 39.7 | 814.0 | 7.04 | 0.18 | n/a | n/a | n/a | 0.77 | 1.10 | 1.71 | 0.53 | 0.57 | 1.17 | 0.53 | 1.63 | 1.5 | 6.5 |
| DV.7.1 | 50.9 | 28.4 | 82.1 | 5.49 | 0.19 | n/a | n/a | n/a | 1.54 | 3.01 | 3.77 | 1.96 | 2.23 | 5.43 | 0.95 | 1.63 | 0.9 | 23.9 |
| BA.4/5 | -27.6 | 60.7 | 3987.2 | 2.37 | 0.33 | 130.80 | 43.05 | 14.74 | 1.54 | 2.11 | 2.87 | 1.30 | 1.58 | 3.54 | 0.94 | 1.48 | 1.5 | 11.7 |
| BQ.1.1 | 2.5 | 53.8 | 1456.9 | 4.82 | 0.26 | 123.00 | 48.46 | 16.57 | 1.34 | 1.69 | 2.19 | 0.54 | 0.49 | 1.38 | 0.81 | 1.39 | 1.8 | 5.7 |
| XBB | 3.6 | 47.5 | 727.9 | 14.30 | 0.20 | 292.70 | 178.10 | 61.29 | 0.40 | 0.63 | 1.13 | 0.50 | 0.71 | 1.41 | 0.32 | 1.45 | 1.0 | 14.7 |
| XBB.1 | 7.4 | 44.7 | 1065.4 | 30.70 | 0.15 | n/a | n/a | n/a | 0.29 | 0.57 | 0.90 | 0.89 | 1.26 | 2.42 | 0.48 | 1.15 | 1.1 | 7.3 |
| XBB.1.16 | 37.6 | 48.3 | 372.5 | 4.69 | 0.11 | n/a | n/a | n/a | 1.08 | 1.59 | 1.94 | 1.32 | 1.18 | 4.27 | 0.85 | 1.67 | 1.1 | 7.2 |
| XBB.1.5 | 24.1 | 45.0 | 475.1 | 3.21 | 0.23 | 182.70 | 61.79 | 22.97 | 1.58 | 2.34 | 3.17 | 0.92 | 1.07 | 2.25 | 0.83 | 1.96 | 1.3 | 10.1 |
| FD.1.1 | 34.4 | 41.1 | 491.1 | 3.60 | 0.09 | n/a | n/a | n/a | 1.39 | 2.51 | 3.55 | 1.06 | 1.09 | 3.05 | 0.86 | 1.79 | 1.0 | 5.7 |
| EG.5.1 | 48.1 | 44.4 | 425.5 | 2.76 | 0.19 | n/a | n/a | n/a | 1.07 | 1.75 | 2.28 | 1.19 | 2.36 | 4.61 | 0.71 | 1.51 | 1.2 | 22.5 |
| HK.3 | 60.9 | 35.4 | 95.9 | 2.39 | 0.25 | n/a | n/a | n/a | 2.10 | 3.12 | 3.54 | 2.00 | 1.83 | 7.44 | 1.10 | 3.29 | 0.9 | 32.1 |
| BA.2.86 | 65.3 | 48.7 | 448.9 | 1.13 | 0.27 | 63.87 | 37.59 | 11.82 | 5.17 | 7.88 | 8.89 | 4.15 | 4.81 | 7.89 | 1.55 | 3.72 | 1.6 | 11.3 |
| JN.1 | 92.4 | 37.1 | 300.8 | 4.62 | 0.15 | n/a | n/a | n/a | 1.66 | 3.40 | 5.05 | 1.15 | 1.47 | 6.11 | 1.27 | 2.44 | 1.5 | 12.3 |

Effect of changes
Decrease Increase

We observed significant differences in the capacity of individuals receiving a fifth dose of bivalent (BA.1 or BA.4/5) mRNA vaccine to recognize and neutralize recent Omicron subvariants (Figure 1B-C). The evolutions of CH.1.1 to DV.7.1, of EG.5.1 to HK.3 and of BA.2.86 to JN.1 led to an improvement in their capacity to evade humoral responses, which could likely be explained by the acquisition of mutations on either S:L455 or S:F456 (30-32, 84, 85). Consistent with other reports, we found that BA.2.86 possesses slightly better escape from plasma-mediated neutralization compared to XBB.1.5, and this was further pronounced with JN.1 (30, 32, 35, 85, 87, 89, 90). These results demonstrate that the high levels of preexisting immune pressure against SARS-CoV-2 Spike glycoprotein is driving its evolution and transmission (48).

By using various approaches, we characterized the ACE2 binding properties of recent Omicron subvariants Spikes. We found that among all Omicron subvariants tested, BA.2.86 was the most susceptible to ACE2-Fc neutralization and exhibited the highest RBD affinity, protomer cooperativity and ACE2 binding at the surface of transfected cells and viral particles (Figure 2-4). These results are in line with previous studies demonstrating the marked improvement in ACE2 binding affinity, explained notably through the acquisition of the S:R403K mutation (33, 88). Furthermore, this increased binding affinity could also be linked to the intrinsic charge properties of the region of ACE2 targeted by the Omicron RBD, which are negatively and positively charged respectively (107-109). Notably, the additional positive charges within the RBD associated with mutations V445H, N460K, N481K and A484K might contribute to its improved ACE2 binding (88). Recently, a BA.2.86 sublineage, JN.1, has shown improved growth rate due to one substitution S:L455S, improving its neutralization escape at the expense of ACE2 binding (32). In this study, we confirm these results by which JN.1 possesses improved immune escape concomitant with its lower ACE2 binding. This further demonstrates the delicate balance existing between neutralization escape and ACE2 affinity in SARS-CoV-2 transmission (5).

It has been suggested that the optimal air temperature for SARS-CoV-2 transmission ranges from 5 to 15°C (110, 111). Within upper airways, lower temperature creates a gradient of temperature from the nasal cavity to the trachea, where it reaches around 33°C (112-114). We previously found that lower temperatures improve RBD-ACE2 interaction and enhance the Spike propensity to sample the ‘’up’’ conformation leading to higher ACE2 binding, fusogenicity and viral replication (52, 55, 56). By priming SARS-CoV-2 Spike, lower temperatures could enhance its ability to bind ACE2 in the upper airways, favoring the initial adsorption (115). In this study, we found that all Omicron subvariants Spikes tested remained sensitive to the impact of low temperature, further enhancing their protomer cooperativity and ACE2 binding at the surface of transfected cells and viral particles (Figure 3-4). More importantly, we found that most Omicron subvariants reached similar levels of binding at 25°C than that of D614G at 4°C at the cell surface and with the RBD, suggesting a lesser reliability on cold temperatures for their improved ACE2 binding. This impact was most notable with DV.7.1, HK.3 and BA.2.86. While multiple factors are involved in SARS-CoV-2 transmission, several lines of evidence demonstrated an association between the climate and increased disease transmission, with lower temperature and humidity being associated with higher COVID-19 incidence (116-124). Whether this is due to ACE2 binding at lower temperature remains to be determined. Notably, the rapid spread of BA.2.86 and expansion of its sublineage JN.1 in North America and in ‘’colder’’ Northern European countries such as the United Kingdom, Denmark, Sweden, Iceland and France raises the intriguing possibility that lower temperature could play a role in transmission through improved ACE2 binding (30, 32, 33, 87, 125, 126).

The Spike glycoprotein requires the adoption of the RBD ‘’up’’ conformation to interact with its receptor ACE2 (44, 45, 127). One parameter affecting the propensity to adopt the ‘’up’’ conformation, and thus ACE2 interaction, is the degree of inter-protomer cooperativity upon ACE2 binding within Spike trimers (61, 128). We and others have demonstrated that SARS-CoV-2 Spike from early variants such as Alpha, Delta and Omicron (BA.1 and BA.4/5) have an improved inter-protomer cooperativity compared to D614G, which was further enhanced at low temperatures (55, 56, 128). Here we show that Omicron evolution continues to follow this trend. We demonstrate a marked improvement in cooperativity among recently emerging Omicron subvariants, with DV.7.1, HK.3, BA.2.86 and JN.1 showing the highest level of cooperativity at 37°C compared to D614G, with these differences being further pronounced at low temperatures (Figure 4). Interestingly, the convergent evolution of emerging Omicron sublineages harboring mutations at either S:L455 or S:F456 illustrates the importance of these residues in transmission (30, 84). Our results show that the acquisition of the ‘’FLip’’ mutations strongly enhanced inter-protomer cooperativity, concomitant with higher ACE2 binding, with DV.7.1 and HK.3 showing a remarked improvement compared to their respective parental lineages (31, 84). Thus, the ongoing surveillance is most likely due to the concomitant improvement in neutralization escape, enhanced protomer cooperativity, and increased ACE2 binding exhibited by the emerging Omicron subvariants, notably facilitated by the ’FLip’ mutations.

We also observed differences in S cleavage to S2 of cells expressing emerging Omicron subvariants Spikes (Figure 5A). In agreement with previous reports, we found that BQ.1.1 and CH.1.1 were processed more efficiently than their respective parental lineages, which could impact their fusogenicity and intrinsic viral pathogenicity (24, 81, 129). We also found that BA.2.86 and JN.1 were more processed than XBB.1.5 and EG.5.1, likely explained by the S:P681R mutation known to enhance Spike processing, fusogenicity and pathogenicity (33, 41, 129, 130). Whether this enhanced processing will affect pathogenicity in humans remains to be known.

We also found a negative correlation between S2 processing and susceptibility to cold inactivation (Figure 5). In contrast to the early Omicron BA.1, recent Omicron subvariants possess enhanced processing while remaining remarkedly sensitive to cold inactivation (102). Interestingly, it was found that susceptibility to cold inactivation may reflect the propensity to sample more ‘’open’’ Spike conformation, thus facilitating conformational transitions (102, 105). These results suggest that, at least for Omicron subvariants, S2 processing could modulate Spike conformation to adopt more ‘’open’’ conformation, rendering them more susceptible to cold inactivation. Interestingly, DV.7.1 and HK.3, harboring the ‘’Flip’’ mutations, had less S2 processing and were less sensitive to cold inactivation compared to their parental lineages, suggesting that ‘’Flip’’ mutations might improve stability through decreased processing, while also strongly enhancing ACE2 binding and the effect of low temperatures. Of note, EG.5.1 was more resistant to cold inactivation than FD.1.1 and XBB.1.5, which could possibly be attributed to its S:Q52H mutation. Interestingly, it was found that EG.5.1 had an increased transmissibility and altered tropism from that of XBB.1.5 (131). Whether this is linked to its higher susceptibility to ACE2-Fc and improved stability remains to be established.

Finally, we also observed that plasma-mediated recognition, neutralization, and ACE2-Fc binding at the surface of cells or pseudoviral particles was associated with the growth rates of emerging Omicron subvariants (Figure 7). The combination of both escape from plasma and ACE2 binding at low temperatures enhanced these associations. These observations further support that ACE2 interaction and immune escape is associated with SARS-CoV-2 transmission and evolution (54, 106). Intriguingly, we observe a higher correlation coefficient for ACE2 binding and virus capture with growth rates at lower temperatures, thus suggesting that temperature modulation of Spike-ACE2 interaction plays a role in viral transmission.

The continued evolution of SARS-CoV-2 requires constant monitoring of its ability to evade immune responses elicited by previous infections and/or vaccination. Our results suggest that the capacity of new emerging variants to interact with ACE2, particularly at low temperatures, is another parameter that deserves to be closely monitored. The growth advantage of XBB.1.5 and its sublineages compared to XBB demonstrates the importance of monitoring ACE2 interaction (28, 29, 40, 44, 98, 106). Although the exact mechanisms through which temperature affects SARS-CoV-2 transmission remain unclear, our findings indicate that Omicron subvariants have undergone mutations that enhance resistance to neutralization by plasma, improve S processing, and increase affinity for ACE2 at both low and high temperatures. Of note, we found that measurement of Spike-ACE2 interaction of viral particles at low temperatures is strongly associated with Omicron subvariants growth rates. Such measures can readily be performed upon the emergence of new variants and could help define which variants have the potential to rapidly expand. In summary, our study underscores the necessity for ongoing surveillance of emerging subvariants and their characteristic mutations, as this information is likely to inform which variants have the potential to become predominant and inform the development of vaccines and other interventions.

## ACKNOWLEDGMENTS

The authors are grateful to the donors who participated in this study. The authors thank the CRCHUM BSL3 and Flow Cytometry Platforms for technical assistance. We gratefully acknowledge the GISAID Initiative and the generous contribution of all data contributors, including the authors, their laboratories that collect the specimens and generated the genetic sequence and metadata on which part of this research is based. Panel A of Figure 1 and Panel A of Figure 2 were prepared using illustrations from BioRender.

M.B. and A.F. conceived the study. M.B., S.D., E.B., A.T., H.M., J.F., M.P., I.L., and C.A. performed, analyzed, and interpreted the experiments. H.M., O.E.F., Y.B., C.B., J.F., M.P., M.C., and A.F. contributed unique reagents. R.P and J.H. performed lineage growth rate estimation analyses. M.B. and A.F. wrote the manuscript with input from others. All authors have read and agreed to the published version of the manuscript.

## DATA AVAILABILITY STATEMENT

Data and reagents are available upon request.

## ETHICS APPROVAL

The study was conducted in accordance with the Declaration of Helsinki in terms of informed consent and approval by an appropriate institutional board. The protocol was approved by the Ethics Committee of CHUM (19.381, approved on 28 February 2022).

## DISCLAIMER

The views expressed in this manuscript are those of the authors and do not reflect the official policy or position of the Uniformed Services University, US Army, the Department of Defense, or the US Government.

## SUPPLEMENTAL METHODS

### Protein expression and purification

FreeStyle 293F cells (Invitrogen, Waltham, MA, USA) were grown in FreeStyle 293F medium (Invitrogen) and transfected with a plasmid coding for soluble ACE2 (sACE2, 1-615), ACE2-Fc (1-615) or the CV3-25 mAbs, using ExpiFectamine 293 transfection reagent, as directed by the manufacturer (Invitrogen). One week later, the cells were pelleted and discarded. Supernatants were filtered using a 0.22 µm filter (Thermo Fisher Scientific). The recombinant sACE2 protein was purified by nickel affinity columns, as directed by the manufacturer (Invitrogen). ACE2-Fc and CV3-25 mAb were purified using protein A affinity column (Cytiva, Marlborough, MA, USA), as directed by the manufacturer. Protein preparations were dialyzed against phosphate-buffered saline (PBS) and stored at -80°C in aliquots until further use. To assess purity, recombinant proteins were loaded on SDS-PAGE gels and stained with Coomassie Blue.

### Flow cytometry analysis of cell-surface staining

Using the standard calcium phosphate method, 10 µg of Spike expressors and 2.5 µg of a green fluorescent protein (GFP) expressor (pIRES2-GFP, Clontech) were transfected into 3 × 10^6^ 293T cells. At 48h post transfection, 293T cells were stained with anti-Spike monoclonal antibodies CV3-25 (5 µg/mL) or using ACE2-Fc (10 µg/mL) for 45 min at 37℃, 25℃ or 4℃. Alternatively, to determine the Hill coefficients, cells were preincubated with increasing concentrations of sACE2 (0 to 166 nM) at 37°C or 4°C. sACE2 binding was detected using a polyclonal goat anti-ACE2 (RND systems, Minneapolis, MN, USA). AlexaFluor-647-conjugated goat anti-human IgG (H+L) Ab (Invitrogen) and AlexaFluor-647-conjugated donkey anti-goat IgG (H+L) Ab (Invitrogen) were used as secondary antibodies to stain cells for 30 min at room temperature. The percentage of transfected cells (GFP+ cells) was determined by gating the living cell population based on viability dye staining (Aqua Vivid, Invitrogen). Samples were acquired on an LSRII cytometer (BD Biosciences, Mississauga, ON, Canada) and data analysis was performed using FlowJo v10.3 (Tree Star, Ashland, OR, USA). Hill coefficient analyses were done using GraphPad Prism version 8.4.2 (GraphPad, San Diego, CA, USA). Values obtained for ACE2-Fc were normalized to CV3-25 signal and represented as a fold over D614G at 37℃.

### Virus neutralization assay

For this, HEK293T cells were transfected with the lentiviral vector pNL4.3 R^-^E^−^ Luc and a plasmid encoding the different S glycoproteins at a ratio of 10:1 to produce SARS-CoV-2 pseudoviruses. Two days post-transfection, cell supernatants were harvested and stored at −80 °C until use. For the neutralization assays, 293T-ACE2 target cells were seeded at a density of 1 × 10^4^ cells/well in 96-well luminometer-compatible tissue culture plates (PerkinElmer, Waltham, MA, USA) 24 h before infection. Pseudoviral particles were incubated with several plasma dilutions (1/50; 1/250; 1/1250; 1/6250; 1/31250), or CV3-25 mAb (from 10µg/mL to 0µg/mL), or ACE2-Fc (from 140µg/mL to 0µg/mL) for 1 h at 37°C and were then added to the target cells followed by incubation for 48 h at 37 °C. The cells were lysed, and luciferase activity was measured as described above. The neutralization half-maximal inhibitory dilution (ID_50_) represents the plasma dilution or the protein concentration (IC_50_) to inhibit 50% of the infection of HEK293T-ACE2 cells by pseudoviruses.

### Biolayer interferometry

Binding kinetics were performed using an Octet RED96e system (ForteBio, Fremont, CA, USA) at different temperatures (35°C, 25°C, 15°C) shaking at 1,000 RPM. Amine Reactive Second Generation (AR2G) biosensors (Sartorius, Göttingen, Germany) were hydrated in water, then activated for 300 s with a solution of 5 mM sulfo-NHS and 10 mM EDC (Sartorius) prior to amine coupling. Either SARS-CoV-2 RBDWT produced in-house or SARS-CoV-2 RBD from several Omicron subvariants (purchased from SinoBiological) were loaded into AR2G biosensor at 12.5 µg/mL at 25°C in 10 mM acetate solution pH 5 for 600 s then quenched into 1 M ethanolamine solution pH 8.5 (Sartorius) for 300 s. Loaded biosensor were placed in 10X kinetics buffer (Sartorius) for 120 s for baseline equilibration. Association of sACE2 (in 10X kinetics buffer) to the different RBD proteins was carried out for 180 s at various concentrations in a two-fold dilution series from 500 nM to 31.25 nM prior to dissociation for 300 s. The data were baseline subtracted prior to fitting, which was performed using a 1:1 binding model and the ForteBio data analysis software. Calculation of on rates (kon), off rates (koff), and affinity constants (KD) was computed using a global fit applied to all data.

---

**Supplemental Figure 1.**
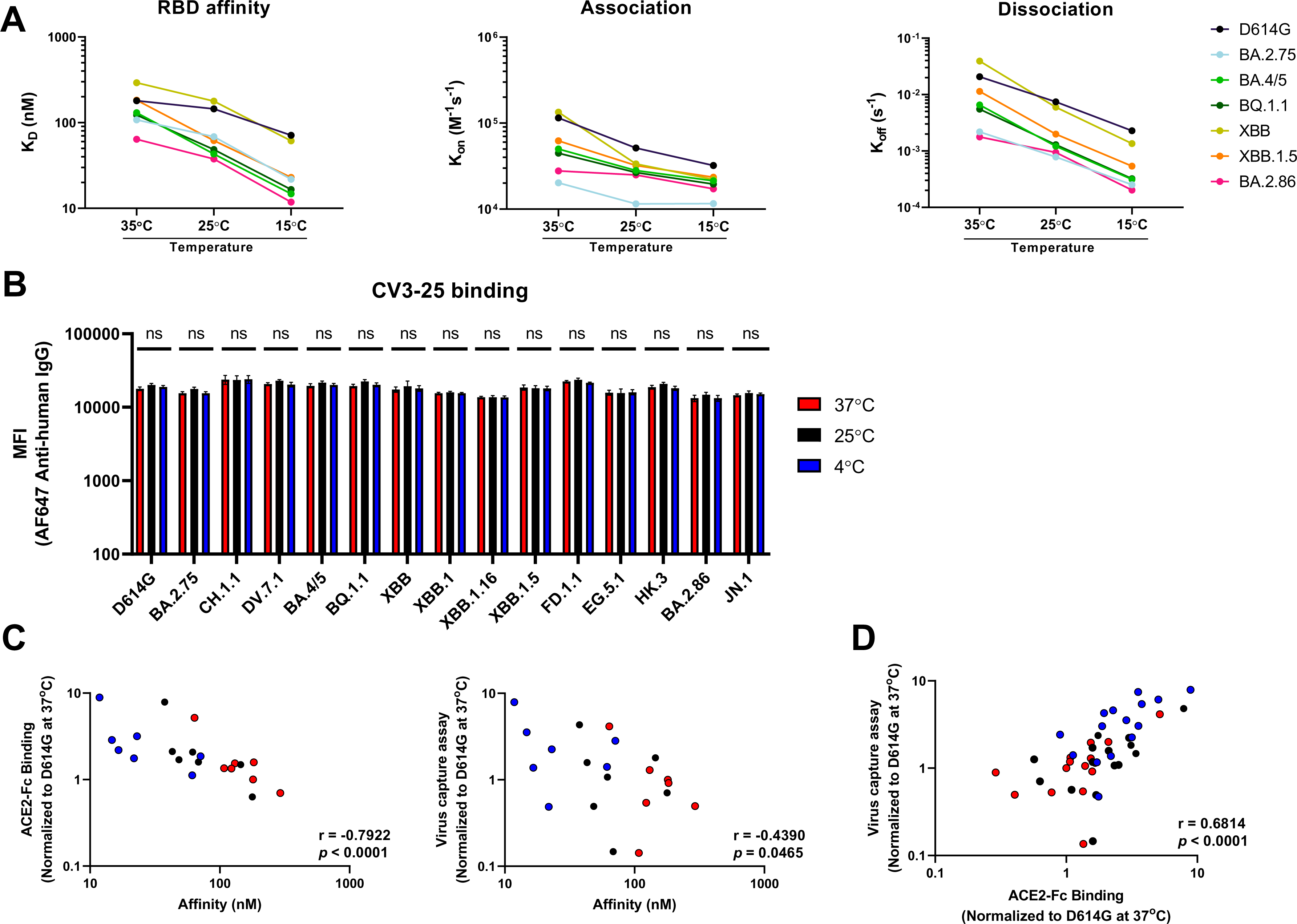
RBD binding kinetics, CV3-25 mAb binding and correlations among techniques at different temperatures. (**A**) Binding kinetics between several RBD (WT or Omicron subvariants) and sACE2 assessed by biolayer interferometry at different temperatures. Graphs represent the affinity constants (K_D_), on rates (K_on_), and off rates (K_off_) values obtained at different temperatures and calculated using a 1:1 binding model. (**B**) Cell surface staining of HEK293T cells expressing full length SARS-CoV-2 Spike glycoproteins from indicated variants (D614G and Omicron subvariants). Temperature-independent CV3-25 mAb is used to quantify the amount of Spike expressed on the cell surface. The graph presents the median fluorescence intensities (MFI). Error bars indicate means ± SEM. These results were obtained in at least three independent experiments. Statistical significance was tested using Mann-Whitney U test (ns, non-significant). (**C-D**) Spearman Rank correlations between RBD binding affinity and cell surface staining or virus capture assay at different temperatures (**C**), and between cell surface staining with virus capture assay at different temperatures (**D**). Panel A, C and D refers to data shown in Figure 3.

